# PRIMATE LIMB CONFORMATION: IMPLICATIONS FOR MECHANICAL ENERGY CONSERVATION*

**DOI:** 10.1101/2021.03.23.436575

**Authors:** Michael P. Catalino

## Abstract

In order to understand the energy implications of primate limb conformation, the biomechanics of swing phase and the vertical movements of limb center of gravity were examined in the arboreal rhesus macaque (*Macaca mulatta*) and the semi-terrestrial long-tailed macaque (*Macaca fascicularis*). The objective was to better understand the potential variation in locomotor adaptations in these two macaque species within different ecological environments. In particular, the objective of this study was to identify the implications that these movements have on the internal energy costs of locomotion. I show that the biomechanics of swing phase for these two species have important effects on their limb center of gravity, although they may have similar limb segment mass distributions. This study suggests that mechanical energy conservation during swing phase is important in mammalian locomotion.

## INTRODUCTION

There is extensive research on animal locomotion (Ahn, Furrow, & Biewener, 2004; Alexander, 1977; Alexander and Maloiy, 1984; Cannon and Leighton, 1994; Cavagna and Kaneko, 1977; Griffin, Main, and Farley, 2004; Kimura, 1992; Kurland et al., 1979; and numerous others), as well was the kinetic characteristics of locomotion for primates and other mammals (Fenn, 1930; Steudel, 1990a, 1990b; Myers and Steudel, 1985; Cavagna and Kaneko, 1977; Griffin et al., 2004; Marsh et al., 2004; Alexander, 1991; Fedak, Heglund, and Taylor, 1982; Heglund, Cavagna, and Taylor, 1982; Heglund, Fedak, Taylor, Cavagna et al., 1982; Taylor, Heglund, McMahon, and Looney, 1980; Manter, 1938; Kimura, Okada, and Ishida, 1979; Pike and Alexander, 2002; Steudel-Numbers, 2003). However, further comparative studies are needed to fully comprehend the complexity of primate locomotion. It is important to understand the means by which primates move as well as the biological and physical factors that make each species unique in their preferred style of locomotion. In an attempt to further comprehend these unique characteristics of locomotion, two Old World monkeys that occupy different niches in nature were compared, the rhesus macaque (*Macaca mulatta*) and the long-tailed macaque (*Macaca fascicularis*). The long-tailed macaque is often considered the most arboreal of the macaques (Kurland, 1973; Wheatley, 1978) while the rhesus macaque can be considered semi-terrestrial since they spend much of their time on the ground or on large diameter tree branches and vines (Dunbar, 1989).

Since these two species are taxonomically closely related, both belonging to the genus *Macaca*, they still share many physical and behavior characteristics, but morphology and other selective pressures may be associated with their divergent evolution to distinct arboreal and semi-terrestrial niches. The mechanics of swing phase and the vertical movements of limb center of gravity were examined to better understand the potential variation in locomotor adaptations between these two macaques in these different ecological environments. In particular, the objective of this study was to identify the implications that these movements have on the internal energy costs of locomotion.

### Effects of gravity on primates

In nonhuman primates gravity plays a more important role than is other mammals, due to the diversity of their locomotor patterns and their variable orientation of body segments in relation to the gravity vector (Preuschoft, 1990). Center of gravity^*^ is defined as the point on which the forces of gravitational acceleration act, and plays a large role in animal locomotion (Preuschoft, 1990). Since all animals can be divided into rigid segments, each segment possesses its own set of physical properties (Manter, 1938) and is subject to the effects of gravity. Many of these properties can be summed to give a representative parameter for the entire body, such as the full body center of gravity. The full body center of gravity determines the load on each limb (Vilensky, 1979). If it is not directly in the center of the area formed by the limb contact points some limbs will hold a disproportionate load compared to the others. Issues of stability can also come into play because the center of gravity needs to be close to the substrate for stability (Schmitt, 1999). Stability can be defined as the vertical line drawn through the center of gravity and it should fall inside the area defined by the points of foot contact (Alexander, unknown). Preuschoft (1990) points out that primates are able to use the earth’s ever-existing source of force in locomotion. This is an important statement that deserves careful consideration in the study of animal locomotion.

### Primate limb design: limb center of gravity implications

For quadrupeds, their gait is traditionally described as symmetrical, where their left and right legs, both fore and hind, commit to a pattern that is a half stride out of phase (Alexander, 2003). In general, primates use a relatively compliant, or bent limb, walking gait when compared to other mammals (Schmitt, 1999). Arboreal primates walking on thin branches have been found to use more compliant gaits for increased stability or due to other selective pressures (Raichlen, 2006; Schmitt, 1999, 2003), and thus arboreal species may have a more compliant walking gait on the ground as well.

If the limbs of animals stayed perfectly straight all the time it could act as a simple natural pendulum, and if every animal had the same length limbs with the same mass and same mass distributions, then the center of gravity would not deviate from pendular motion when the animal moves. However, this is not the case. The primate limb is comprised of three segments: proximal, middle, and distal (Manter, 1938). Each segment has a mass and length and therefore a mass distribution. Limb length, mass, and mass distribution are highly variable across animal species (Fedak et al, 1982; Vilensky, 1979; Reynolds, 1985a; Raichlen, 2006) and primates in particular have longer limbs than typical mammals (Alexander, 1985). As the limb conformation changes (Figure 3), the center of gravity moves away from the limb and can change location in three dimensions. Therefore, changes in limb properties and movements during walking affect the movements of the actual limb center of gravity.

During stance phase for example, the center of gravity can partly determine the direction and magnitude of substrate reaction forces which create bending moments on limb joints (Schmitt, 1999).

Intuitively, the forces generated by the downward acceleration of the body’s center of gravity are transmitted to the ground through the limbs. Newton’s third law states that for every action there is an equal and opposite reaction. This substrate reaction force has been shown to be the main cause of the bending of limbs due to the moments that are created by the joint angles (Schmitt, 1999, 2003).

Disregarding the center of gravity of the whole body and focusing on the center of gravity of each limb, we can move forward from stance phase and look at the dynamics of swing phase. Instead of the body’s center of gravity acting through the limb, the limb is changing its center of gravity with relation to that of the body’s and even using the body’s center of gravity to accelerate the limb forward during this aerial phase. When the muscles act on the limb, they are actually shortening both attachments, but the moment of inertia of the entire body is large compared to the limb, and is more difficult to move, causing the limb to accelerate passed the body during swing phase.

### Quadrupedal gait: Walking in primates

Walking is the style of gait that animals use to maximize the efficiency of locomotion (Alexander, 2003). Therefore, for mammals that move around by primarily using their legs, getting from one place to another is usually accomplished by a walking gait (Griffin, et al., 2004). A walking gait is characterized as having at least one limb in contact with the substrate throughout each stride (Alexander, 2003). Duty factor reveals the amount of time each limb spends in contact with the ground, doing work by exerting forces, as a percent of the total stride duration (Griffin, et al., 2004), and walking requires a duty factor that is greater than 0.5 (Alexander, 2003). Other types of gait exists for animals, and for quadrupedal mammals, these can include galloping, running, and hopping (Alexander, 2003). Since most animals spend so much of their time walking, it is important to consider all aspects of animal walking. Animal walking involves the limbs acting as support for the body during repeated, cyclic motions. Each stride of an animal can be broken up into the stance phase and the swing phase, where the swing phase is a period of high velocity and the stance phase is a period of low velocity (Manter, 1938). During this time, for primates, the limb does not remain totally straight, and in fact, each joint goes through many angular excursions throughout the stride (Kimura, Okada, and Ishida, 1979; Pike and Alexander, 2002; Hildebrand and Hurley, 1985). These changes in limb conformation are important, because if we are going to treat the limb as separate rigid segments (Manter, 1938), than the dynamics of the limb will be affected by the movements of each segment.

### Energetics of animal locomotion

Current research reveals that deviation from preferred style of locomotion does cause changes in the energetic cost (Nakatsukasa, et al., 2004). In addition, the mechanical power trade-off mechanism of locomotion is affected by muscle mass distribution, and varies across mammalian taxa (Raichlen, 2006). The energetic cost of locomotion is also affected by the center of gravity. Modeling the limb as a natural pendulum (Hildebrand and Hurley, 1985, Myers and Steudel, 1997), it has a swing period was found to coincide significantly with the stride durations of animals walking at moderate speeds (Hildebrand and Hurley, 1985). He suggests longer limb length is an adaptation that favors economic walking over long distances. By modeling a natural pendulum the limb would swing naturally and therefore minimal energy would be expended.

Efficient runners are found to have more proximal limb center of mass (Taylor et al, 1974). This suggests, that for limb design, even minor changes in limb mass and mass distribution may have a relatively large affect on energy consumption (Taylor, Shkolnik, Dmi’el, Baharav, and Borut, 1974; Raichlen, 2006). The center of gravity movements and velocity can be used to calculate kinetic energy (KE = 1/2mv^2^), and the height of the center of gravity can be used to calculate potential energy (PE = mgh), where g is the acceleration due to gravity on the earth. Using angular changes in motion, rotational energy can also be calculated. Humans (Cavagna et al, 1982) and dogs (Griffin et al, 2004) use an inverted pendulum to achieve energetic transfer of KE and PE when walking efficiently. Energy is vulnerable to changes in limb mass distribution. Adding mass close to the body’s center of gravity results in a significantly lower increase in energy compared to loading the mass distally (Myers and Steudel, 1985). Center of gravity is also a major component when calculating inertia, which measures resistance to changing motion (Tipler, 1986).

### Primate limb conformation and energy consumption

Since energetic efficiency is important to reduce energetic costs of foraging, the movements of the limb center of gravity may affect the work required for swing phase and therefore the energy consumed during this portion of the stride. It has already been suggested that the location of the limb’s center of gravity affects the energetic costs of swing phase (Taylor et al, 1982; Fedak et al 1982; Raichlen, 2004, 2006; Larson, Schmitt, Lemelin, and Hamrick, 2000). The acceleration of the limb center of gravity relative to the body center of gravity requires energy (Fedak et al, 1982) and is often considered a major component of internal energetic cost (Fedak et al, 1982; Manter, 1938; Steudel, 1990b). Although the mass of each segment decreases as you move distally, the distance the distal segments travel during swing phase is greater and therefore they have larger velocities and larger fluctuations in KE (Fedak et al, 1982). Fedak et al (1982) actually found that kinetic energy seems to increase at about the 1.55 power of speed and not between the square and cube or running speed, as expected, since kinetic energy is proportional to the square of velocity. This result comprised of a propulsive stroke, which is at the end of stance phase, increasing its kinetic energy as the 2.2 power of speed. This falls remarkably close to the estimated cost to increase velocity. However, they also found that during the recovery stroke, which is considered the aerial phase or swing phase, the kinetic energy increased as the 1.1 power of speed. They suggest that the path of recovery stroke appears to minimize increases in limb velocity at high speeds by greater joint flexion, bringing the limb center of gravity closer to the body. To this regard, it is expected to see that the recovery stroke becomes longer at high speeds (Fedak et al, 1982), since it would help decrease the total energetic cost of increasing speed.

Interestingly, Raichlen (2006) pointed out that if the older individuals had the limb mass distributions of juveniles, but did not reduce their stride frequencies, they would have increased their internal work by approximately 20% at a given velocity. In other words, younger individuals save approximately 20% of the mechanical power they would otherwise have had to use to move their limbs relative to their body. Although baboons are mainly terrestrial, juvenile baboons have many features similar to more arboreal primates, such as a more distal limb mass distribution (Raichlen, 2004; Preuschoft and Gunther, 1994). Infant baboons showed lower internal energy and higher external energy consumption when compared with non-primates (Raichlen, 2006), which explains why the total energy may appear similar without breaking it up into the internal and external components. Low stride frequencies that are associated with low internal energy cost, and long strides are associated with high external energy costs (Raichlen, 2006).

If energetic costs weigh heavily on a species as a selective pressure we would expect animals to adapt to be the most efficient in every way. Moving requires energy and therefore, getting from place to place to find food, shelter or escape predation is when most of an animal’s energy is expended. The setting in which the animal spends most of its time will have a great influence on its energy consumption and the role of energetic in locomotor progression (Rodman, 1979). Medium and large sized terrestrial quadrupeds and bipeds have figured out how to use stiff legged walking to convert potential and kinetic energy using the inverse pendulum, conserving about 70% of their energy when walking at certain velocities (Heglund et al, 1982). Arboreal quadrupeds, especially non-human primates, have to deal with an unpredictable environment, small branches and unsteady surfaces (Rollinson and Martin, 1981). In order to reduce branch sway, which would cause increased energy expenditure, the arborealist must decrease its substrate reaction forces produced during stance phase and propulsion in each step (Schmitt, 1999; Larson et al, 2000). Schmitt (2003) suggests that terrestrial primates may be limited in arm protraction that would allow them to increase compliance, reducing their forces applied to the substrate. To accomplish lower substrate reaction forces, primates use long low strides and increased angular excursions (Larson et al, 2000). Walking compliantly may reduce peak vertical forces (Schmitt, 1999; Larson et al, 2000), the longer the limb is in contact with the ground the greater sum of the force is that is exerted over the time of each stride. Therefore, the external work that is done on the limb in propelling the animal’s center of gravity forward increases. As Raichlen (2006) points out, an increase is external work may be associated with a decrease in internal work in order to keep the total work done constant during each stride. It has been suggested that animals bend their limbs to change limb center of gravity, bringing it closer to their body during swing phase (Fedak et al, 1982; Marsh et al, 2004). This is thought to reduce energetic costs of locomotion (Fedak et al, 1982). Since primates have already adapted a compliant gait to lower their peak substrate reaction forces (Schmitt, 1999; Larson et al, 2000), it is logical to look elsewhere for a mechanism that decreases internal energy costs. Since compliance refers to a style adapted during stance phase, swing phase may be the key to lowering internal energy expenditures.

### Components of energetic cost during animal locomotion

The stance phase involves ground reaction forces to propel the body center of gravity forward and therefore has received much attention in the field of animal locomotion (Schmitt, 1999, 2003; Kimura, 1979; Fedak et al, 1982; Heglund et al, 1982). Some say that swing phase contributes so little to the total energy consumption that it is irrelevant (Taylor, 1980), but recent studies have shown that swing phase consumes 26% of the total energy (Marsh et al, 2004). Although it is less studied, swing phase is important in quadrupedal locomotion primarily due to its similarity to the motion of a natural pendulum (Myers and Steudel, 1997; Raichlen, 2004) and is therefore subject to the laws of physics.

Since kinetic energy should increase as the square of velocity, the swing phase becomes increasingly important at higher speeds (Manter, 1938). Fedak et al (1982) found that the increase in the kinetic energy of the limbs relative to the body at higher speeds is almost constant for the ostrich and horse. In addition, it has been suggested that animals use enhanced non-metabolic means to power internal work when moving at higher speeds (Steudel, 1990b). The energy storage and use during running has been focused on elastic storage in ligaments and tendons, acting like a loaded spring during impact (Biewener, 2006). The path of the swing phase also seems to reduce energetic costs at high speeds by bringing the limb center of mass closer to the body (Fedak et al, 1982). If elastic storage is less likely when walking do to the lower force exerted on the ground, the path of the limb center of gravity during swing phase seems to be the best mechanism available for decreasing internal energy costs.

If energy conservation is acting as a strong selective pressure in primates, and gravity plays such a prominent role in energetic costs, it is likely that these animals have adapted locomotor characteristics that exploit these physical limitations. In this study, the swing phases of two closely related primate species will be carefully examined for differences that could have implications in internal energy conservation. The focus of this work is to see the extent to which limb conformation affects the position of the limb center of gravity during swing phase, and if the magnitude of the results differ between the arboreal and terrestrial sample. Arboreal primates are thought to have greater distal limb segment mass for greater grasping ability compared to more terrestrial primates (Preuschoft and Gunther, 1994), and must prevent branch sway when walking on thin branches (Rollinson and Martin, 1981), so I hypothesized that the *M. fascicularis* when compared to the *M. mulatta* will have:

1. less shoulder and hip retraction,
2. greater elbow and knee joint flexion and,
3. larger vertical changes in limb center of gravity during swing phase.

## MATERIALS AND METHODS

### Sample

The mechanics of locomotion were compared across two species of Cercopithecine primates walking on simulated terrestrial substrates in The Locomotion Lab at Duke University. The sample of primate species included one species that can be considered semi-terrestrial, the rhesus macaque (*M. mulatta*) (Dunbar, 1989) and the other, mainly arboreal long-tailed macaque (*M. fascicularis*) (Schmitt, 1999, 2003). Terrestrial activity is dominant for all ages in *M. mulatta* (Wells and Turnquist, 2001), and the *M. fascicularis* spend the majority of their time in the main canopy of forests (Cannon and Leighton, 1994). The *M. fascicularis* are about 50% smaller in body mass than the *M. mulatta*. Female *M. fascicularis* weigh approximately 3.5 kilograms and males about 5.3 kilograms (Fleagle, 1999). Both species have an intermembral index of about 93, which is a ratio of forelimb length over hindlimb length. So, although they are taxonomically similar, the variation across species allows their mechanics of locomotion to be explored and compared in detail with implications for adaptations to arboreal or terrestrial habitat. We focused on the swing phase of the stride cycle, measuring joint angles and using them to follow the vertical movements of the limb’s center of gravity. In addition to the data collected on live primate locomotion, segment center of gravities were conducted at the University of Illinois on an adult long-tailed macaque donated by the Southwest Foundation through Steven Leigh and John Polk, PhD from the University of Illinois and Department of Anthropology.

### Limb segment center of mass

The data were collected from a cadaver obtained from the Southwest Foundation. The protocol was completed for both the fore and hind limbs in their entirety as well as individual limb segments appropriately named the proximal, mid, and distal limb. The protocol for collecting center of gravity and limb segment parameters was similar to the methods in Myers and Steudel (1997) and Vilensky (1979).

The dismemberment was preceded by preliminary limb length measurements. The total limb length (TL) was measured while holding limbs in straightened position. The forelimb TL was measured from the acromion process of the scapula and the greater tubercle of the humerus to the tip of the fore paw. The hindlimb TL was measured from the ischial tuberosity and the greater trochanter of the femur to the tip of the hind paw. The dismemberment removed the forelimbs excluding the scapula, since it was not believed to be involved in the swing phase. (Myers and Steudel, 1997).

To determine where to cut, the gleno-humeral and hip joints were located using palpation. The limbs were straightened and tied to a board to be frozen (limb mass loss due to freezing was negligible according to Myers and Steudel, 1997) and the post freezing limb mass (LM) was used in all calculations. After freezing the following measurements were taken for the forelimb (FL) and hindlimb (HL): distal segment length (DL) for the forelimb – pisiform to the tip of paw and for the hind limb – calcaneum to the tip of the paw, mid + distal segment length (MDL) again for the forelimb – olecranon process to tip of paw and hind limb – intercondylar tubercle of tibia to the tip of paw, midjoint circumference (MJC) for the forelimb – just superior to the olecranon process and hind limb – between the patella and the tuberosity of the tibia, distal joint circumference (DJC) for the forelimb – just superior to the styloid process of the ulna and radius and hind limb – just superior to the calcaneum, proximal leg segment (PL) = TL – MDL, midsegment length (ML) = MDL – DL, and proximal + midsegment length (PML) = PL + ML.

In order to determine the center of mass (CM), each limb was placed on a lightweight aluminum tray and balanced on its fulcrum (Figure 1). The location of the center of mass was calculated relative to the most proximal joint using the system balanced in equilibrium and calculating the sum of torque about the wooden dowel. The limb segment weight and center of gravity as well as the weight and center of gravity of the tray cause opposite and equal torques about the wooden dowel (see Appendix 1).

**Figure 1:**
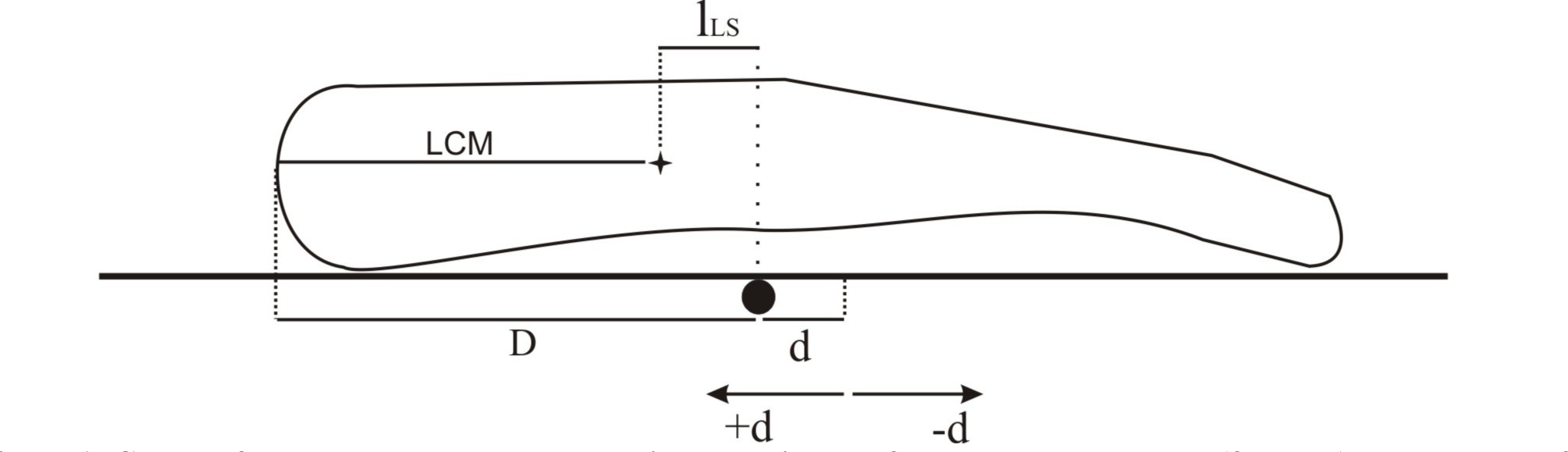
Center of Mass measurements. d describes the distance from the wooden dowel (fulcrum) to the center of mass of the tray, for each trial d could be positive or negative depending on where the tray balanced on the fulcrum . lLS is the moment arm created by the limb center of mass (LCM). The moment arm created by the center of mass of the try is just d. See **Appendix 1** for details on calculating the LCM using summation of torques.

Using summation of torques instead to find the limb center of mass (LCM) instead of moving the limb on the metal tray, as was done in Myers and Steudel (1997), may reduce error caused by friction on the wooden dowel.

### Swing phase mechanics and limb conformation

To analyze the motion of swing phase and how limb conformation affects the vertical displacement of the center of gravity, we analyzed between video taken at the Animal Locomotion Lab as well as previously recorded video data (Schmitt, 1995). Videos of *M. fascicularis* “Jones”, and *M. mulatta* “Jane” were taken using a high speed 60fps video camera. Anterior-posterior as well as medial-lateral video was taken but only the medial-lateral video was analyzed. Successful runs were considered those where one complete swing phase for the fore-and hind limb was completed within the viewing window. Each successful step was recorded to DVD and clipped for analysis using VirtualDubMod.

Once the single step was isolated, the number of frames from lift off to touch down was recorded. In order to control for speed, only steps that were within one 2/60^ths^ of a second (or two frames) of each other were kept for analysis. The video clip was then deinterlaced in order to maintain high resolution during playback in slow motion. I used VirtualDubMod and the “GunnerThalin” deinterlacing filter. Once the video was filtered, it was digitized using MatLab and DLTdataviewer (Hendrick, 2008). I digitized the forelimb and hindlimb facing the camera because this assured the most accurate placement of joint markers. The points digitized were the proximal joint (shoulder or hip), middle joint (elbow or knee), and distal joint (wrist or ankle). We treated the distal limb segment (hand or foot) as a point mass at the end of the middle segment (forearm or thigh), which is an accepted method of study (Raichlen, 2006). Before running the data coordinates through MatLab to obtain joint angles we had to create and horizontal point in order to calculate the shoulder angle. This horizontal point extended cranially for the hip and caudally for the shoulder. We chose this direction so that MatLab would measure the retraction angle with respect to the horizontal. All data points were analyzed using MatLab <RUNJOINT> and <JOINTANGLE>. The resulting data were joint angles throughout the swing phase, one angle for each the proximal and middle joints for each frame. These joint angles and coordinates of joints in space were then used to calculate the vertical movements of the bent limb center of gravity throughout swing phase.

### Bent limb center of gravity

The bent limb center of gravity was found by first analyzing the changes in proximal (θS) and middle joint (θE) angles throughout swing phase as mentioned above. For each frame, the bent limb center of gravity (BLCg) was calculated based on the equation from Appendix 2:

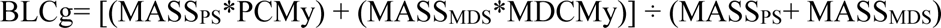

The vertical coordinates of the bent limb center of gravity were calculated for each frame throughout swing phase for each step.

### Manipulation of the center of mass

A manipulation of the limb center of mass location of each of the limb segments to test the sensitivity of the bent limb center of gravity to changes in limb segment center of mass.

In order to test the this effect that limb mass distribution may have on the movements of the bent limb center of mass during the swing phase, we recalculated the vertical changes in bent limb center of gravity for our *M. fascicularis* subject using the limb segment center of mass data found by Vilensky (1979) for *M. mulatta*. Since the limb segment center of mass in Vilensky (1979) are given as dimensionless values (percent total limb length) we did not have to scale the data between individuals. The limb segment mass and joint flexion and retraction angles were kept the same, so that the only difference was the location of the center of mass on each limb segment.

### Data analysis

The number of steps analyzed ranged from n=5 to n=15 for each parameter (Table 1). Peak vertical values of the limb center of gravity were recorded for the two species for each step for both the forelimb and the hindlimb. In order to control for different starting values, the y value that corresponds to the first frame of swing phase is considered the zero line, and all vertical deviations from that line were calculated. Therefore, the peak vertical value describes the maximum height the limb center of gravity reached above this zero line drawn at the initiation of swing phase. Maximum and minimum hip and shoulder retraction, and joint angular excursion was also recorded for each step. Finally, maximum and minimum elbow and knee flexion angle, and joint angular excursion were recorded and compared. All variables were tested for significance across species using a non-parametric two sample Mann-Whitney U test, all significant differences are tentative given the relatively small sample size.

**Table 1:**
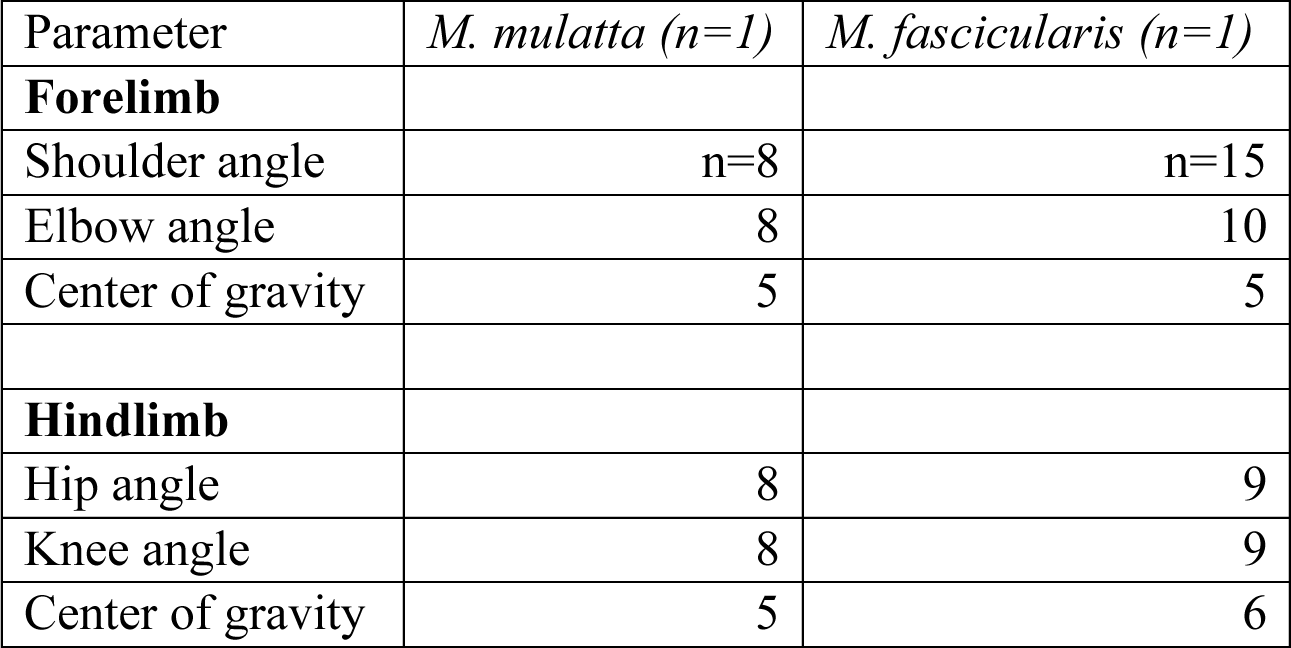
Description of sample and the number of step analyzed for each parameter.

| Parameter | *M. mulatta* (n=1) | *M. fascicularis* (n=1) |
| --- | --- | --- |
| **Forelimb** |  |  |
| Shoulder angle | n=8 | n=15 |
| Elbow angle | 8 | 10 |
| Center of gravity | 5 | 5 |
| **Hindlimb** |  |  |
| Hip angle | 8 | 9 |
| Knee angle | 8 | 9 |
| Center of gravity | 5 | 6 |

**Figure 2:**
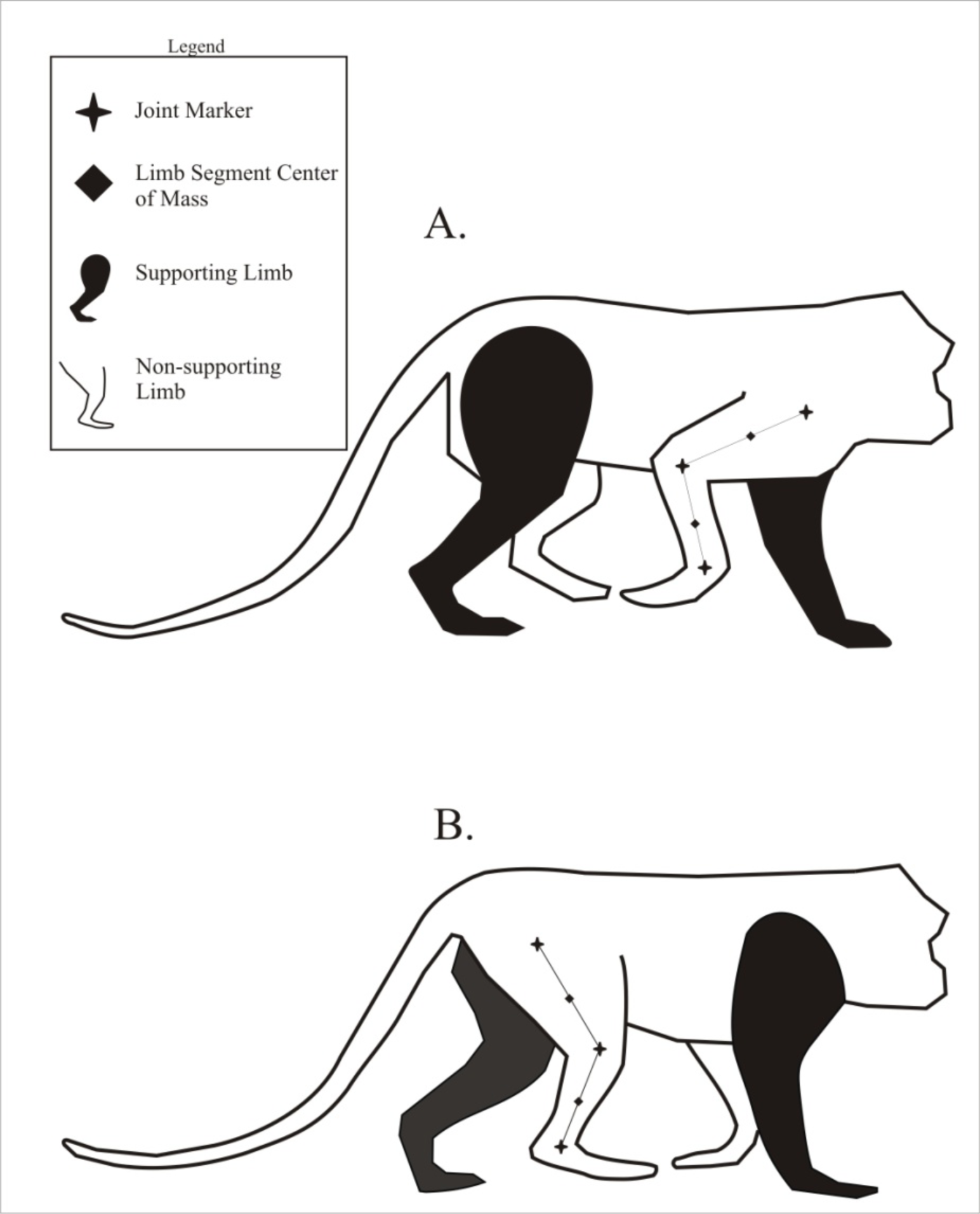
The joints that were digitized using MatLab DLTdataviewer are shown as stars. The limb segment center of mass (diamond) was calculated after using the equations from Appendix 2.

**Figure 3:**
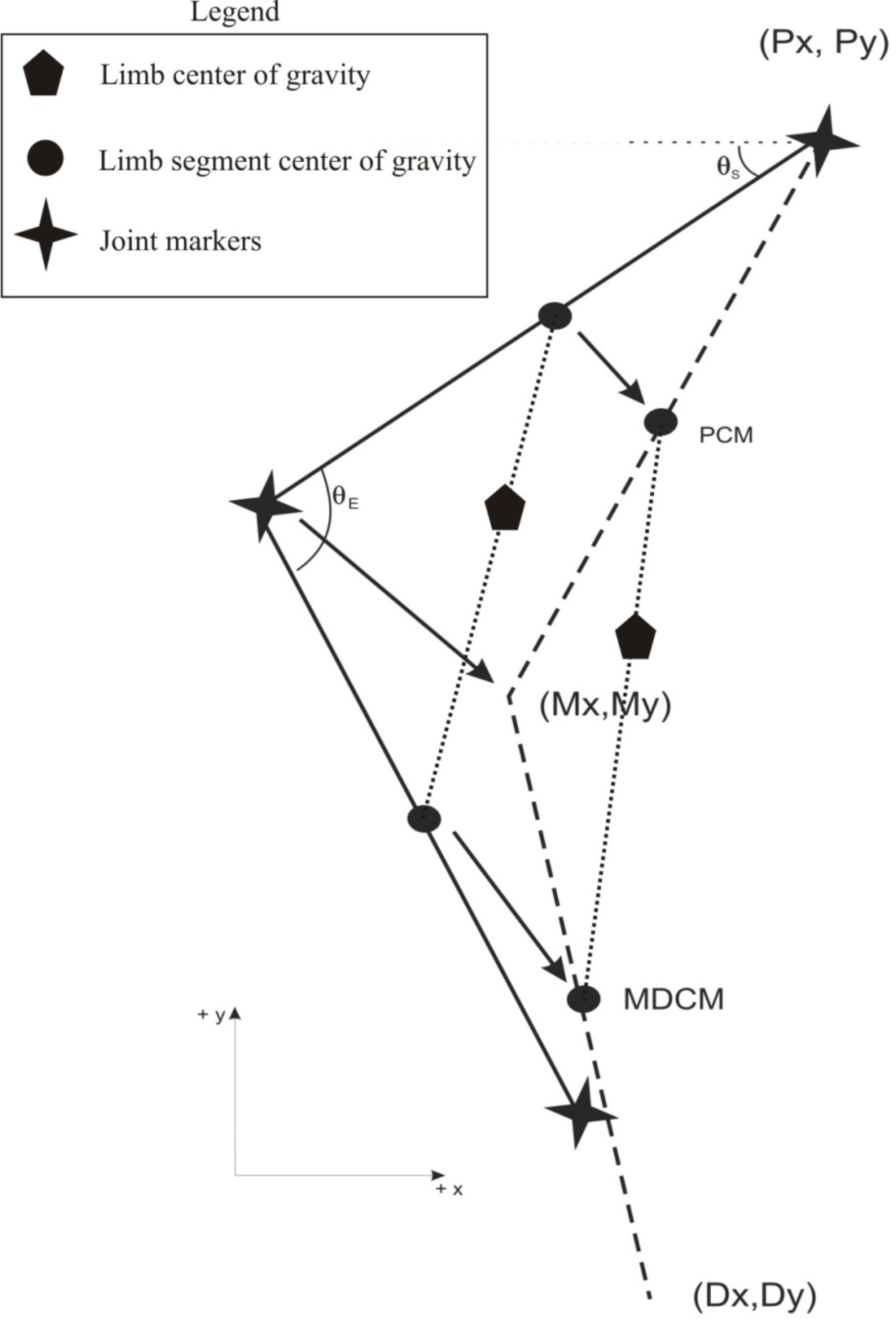
Data points obtained from MatLab DLTdataviewer. The most proximal joint coordinates are (Px,Py), the middle joint coordinates are (Mx,My), and the distal joint coordinates are (Dx,Dy). PCM is the proximal center of mass of the proximal limb segment and MDCM is the mid-distal center of mass of the middle limb segment when the distal limb segment is treated as a point mass at the distal end of the middle segment. θ E is the elbow angle taken from the position of the three mentioned point. θ S is the shoulder angle taken from the horizon line. The limb Cg lies on the line between the two segment center of masses (Tipler, 1986) and can be calculated using the equations in Appendix 2.

**Figure 4:**
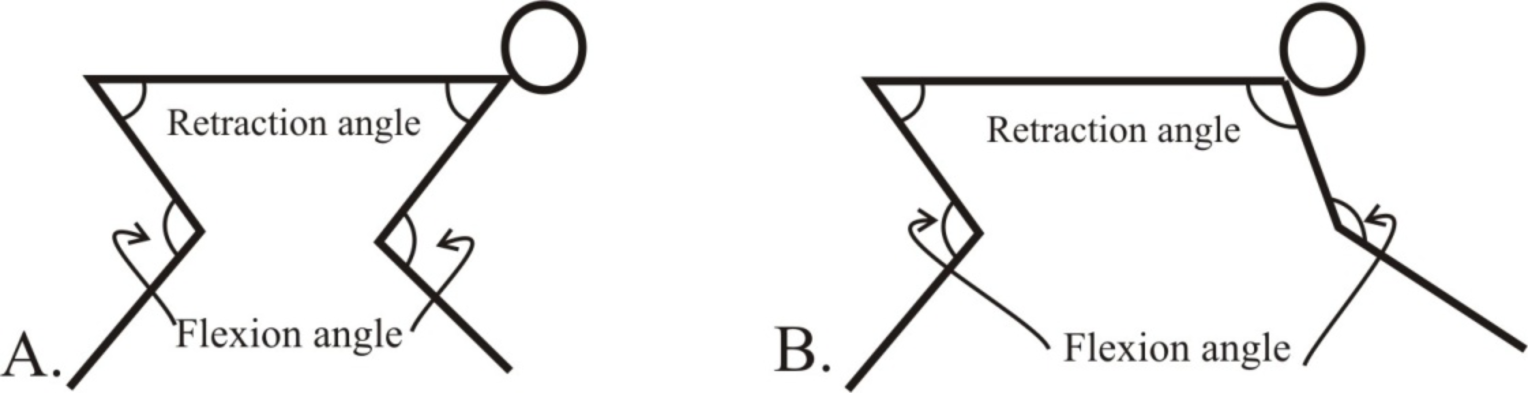
Retraction and flexion angle orientation defined for the hip, shoulder, knee, and elbow of a quadruped for this study. Image B depicts the forelimb from image A protracted, or at a larger than 90 degree retraction angle^1^.

## RESULTS

Recent studies have suggested that limb conformation during swing phase can cause changes in the location of the limb center of gravity relative to the body (Fedak et al, 1982; Raichlen, 2004, 2006; Myers and Steudel, 1985). The data support this and suggests that elbow and knee flexion have the most influence on the vertical peak of the limb center of gravity during swing phase. The *M. fascicularis* showed a significantly higher peak vertical change in both the forelimb and hindlimb center of gravity, when compared to the *M. mulatta*. Hip retraction showed no significant differences between the two individuals. Although maximum retraction angle showed no significant difference, minimum retraction angle was significantly smaller (p = 0.017) and shoulder retraction angular excursion was significantly larger (p = 0.041) for *M. fascicularis.* The maximum knee flexion angle was significantly smaller (p = .039) for the *M. fascicularis*, while the minimum knee flexion and knee angular excursion showed no significant differences. Maximum elbow flexion angle was significantly smaller (p < 0.001) for the *M. fascicularis*, and elbow angular excursion was significantly larger (p < 0.001), while minimum elbow flexion showed no significant differences. The limb center of gravity showed significantly greater vertical peaks for both the forelimb (p = 0.012) and the hindlimb (p = 0.008) in *M. fascicularis*.

### Manipulated center of mass

The resulting changes in center of mass fluctuations caused by a more distal center of mass were still significantly different for the forelimb (p = .012) and for the hindlimb (p = .008) between the individuals (Table 4). This was expected because a more distal limb segment center of mass will have greater vertical fluctuations than a more proximal one for a given change in angle of the joint.

**Figure 5:**
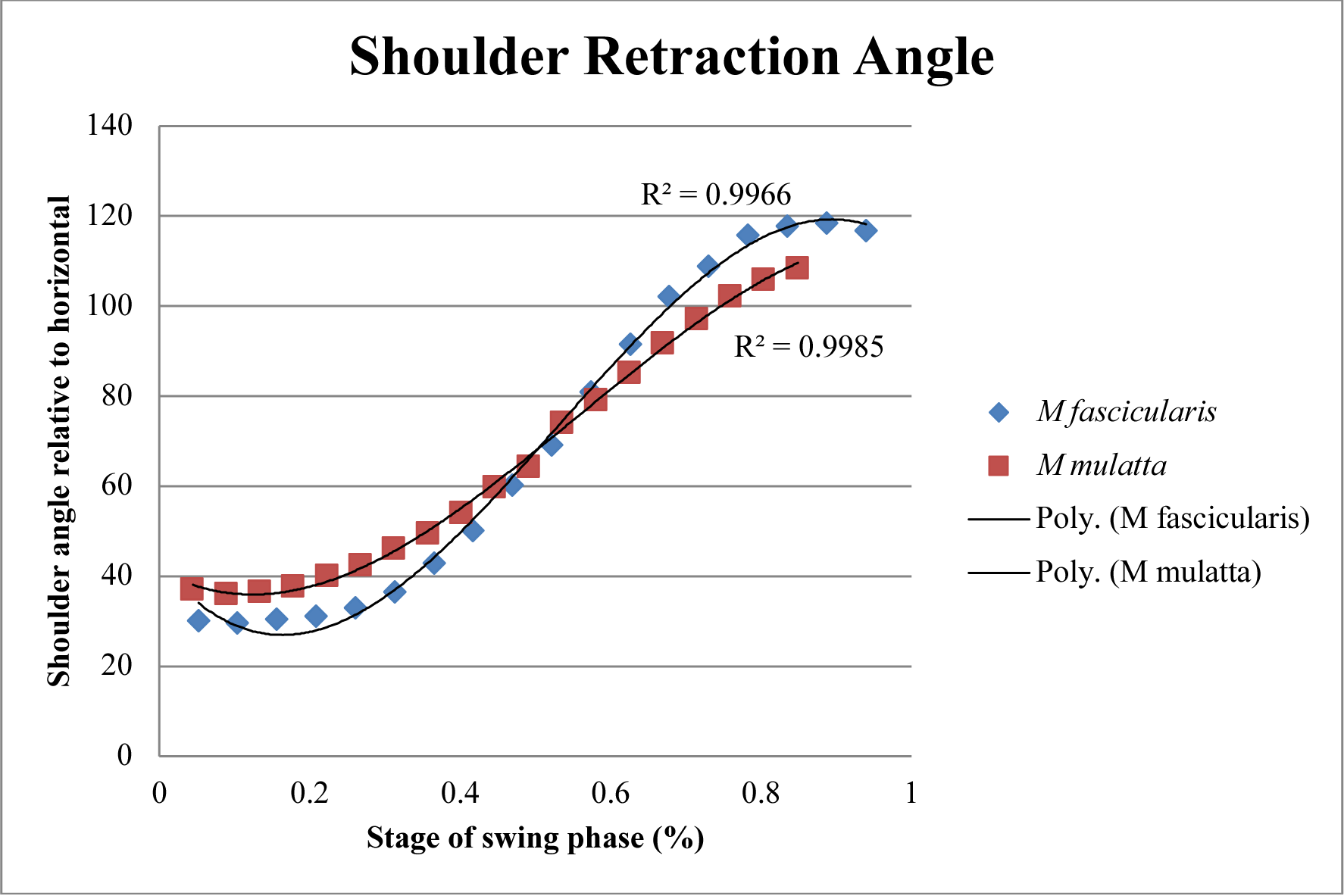

**Figure 6:**
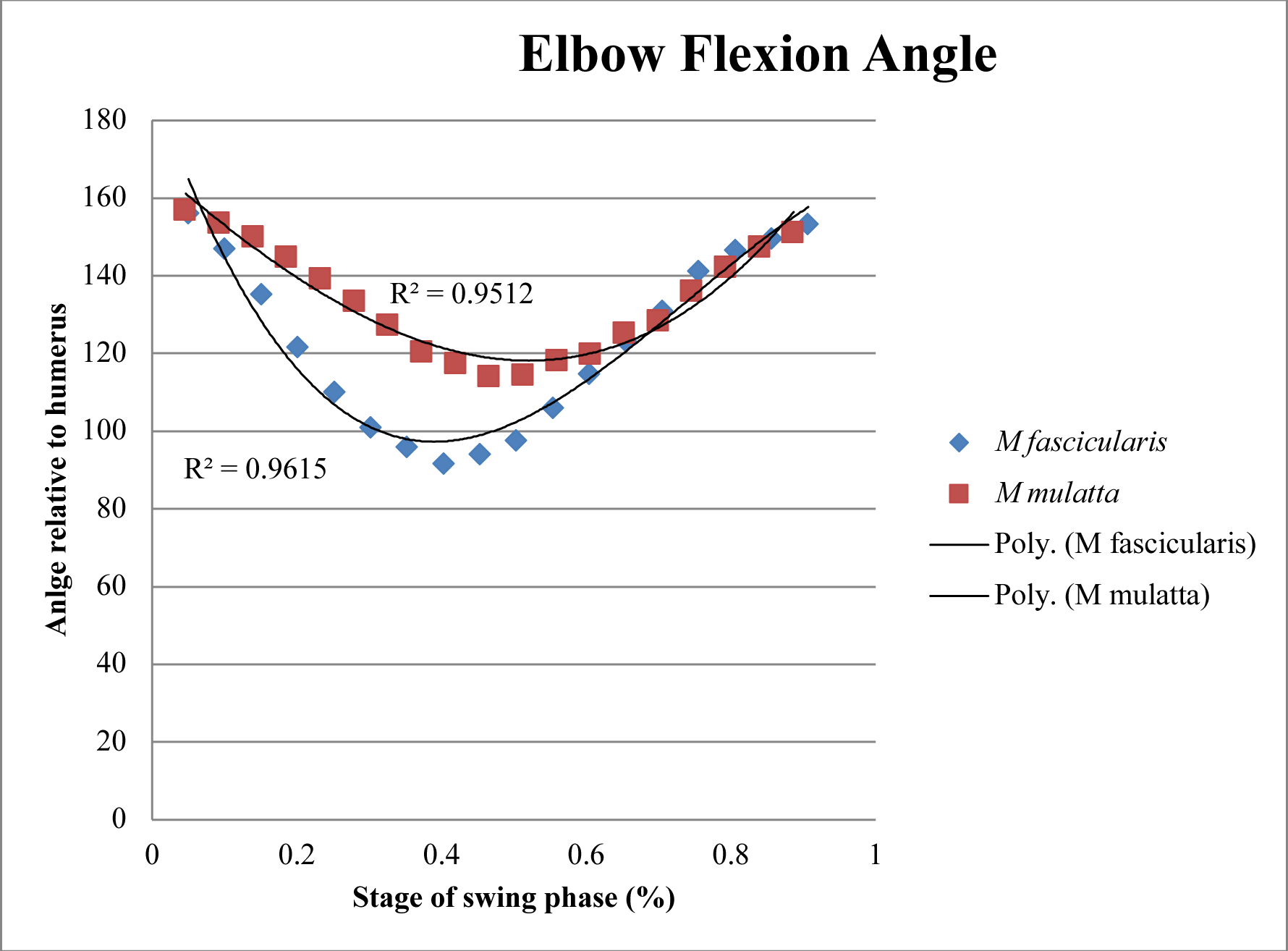

**Figure 7:**
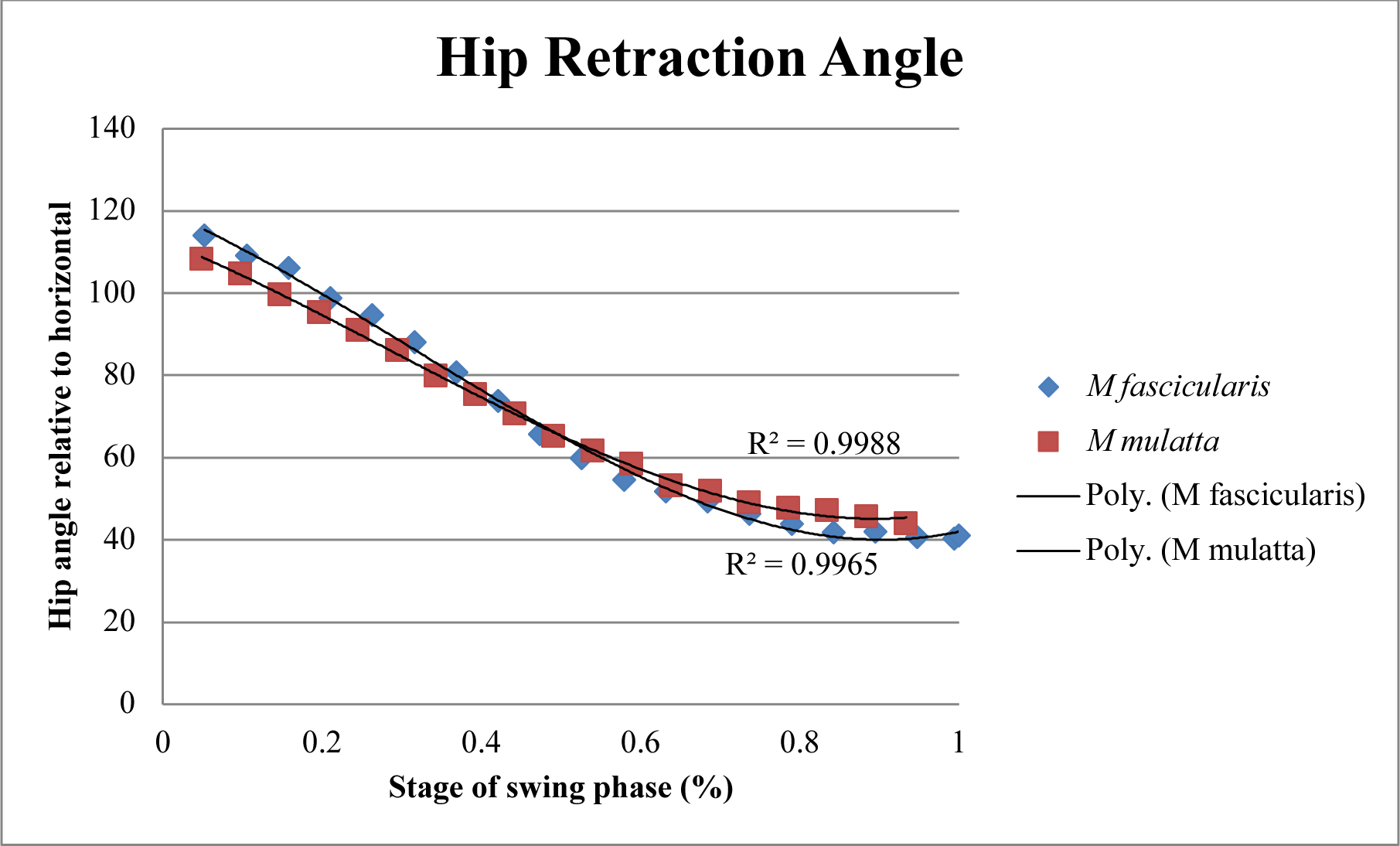

**Figure 8:**
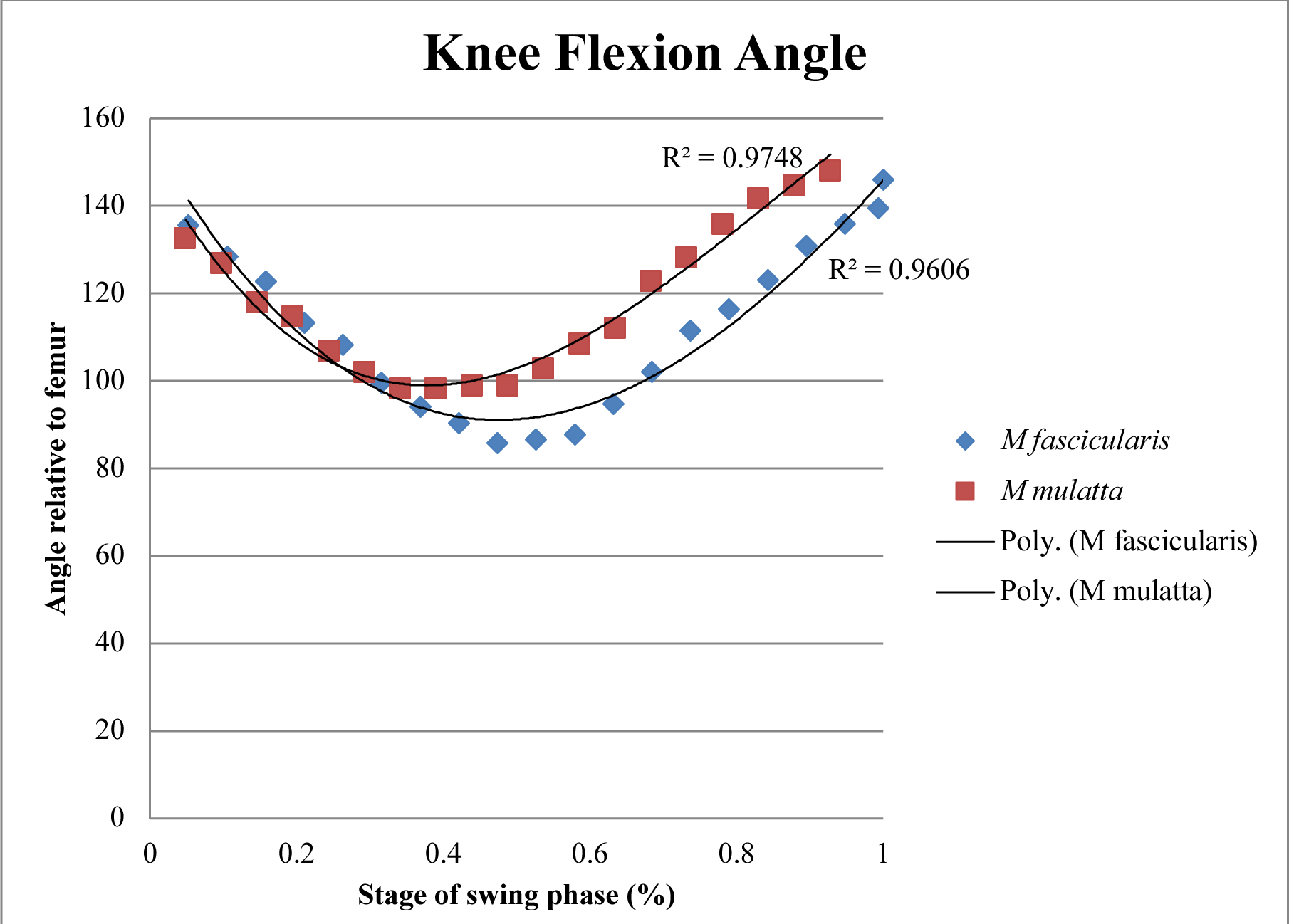

**Figure 9:**
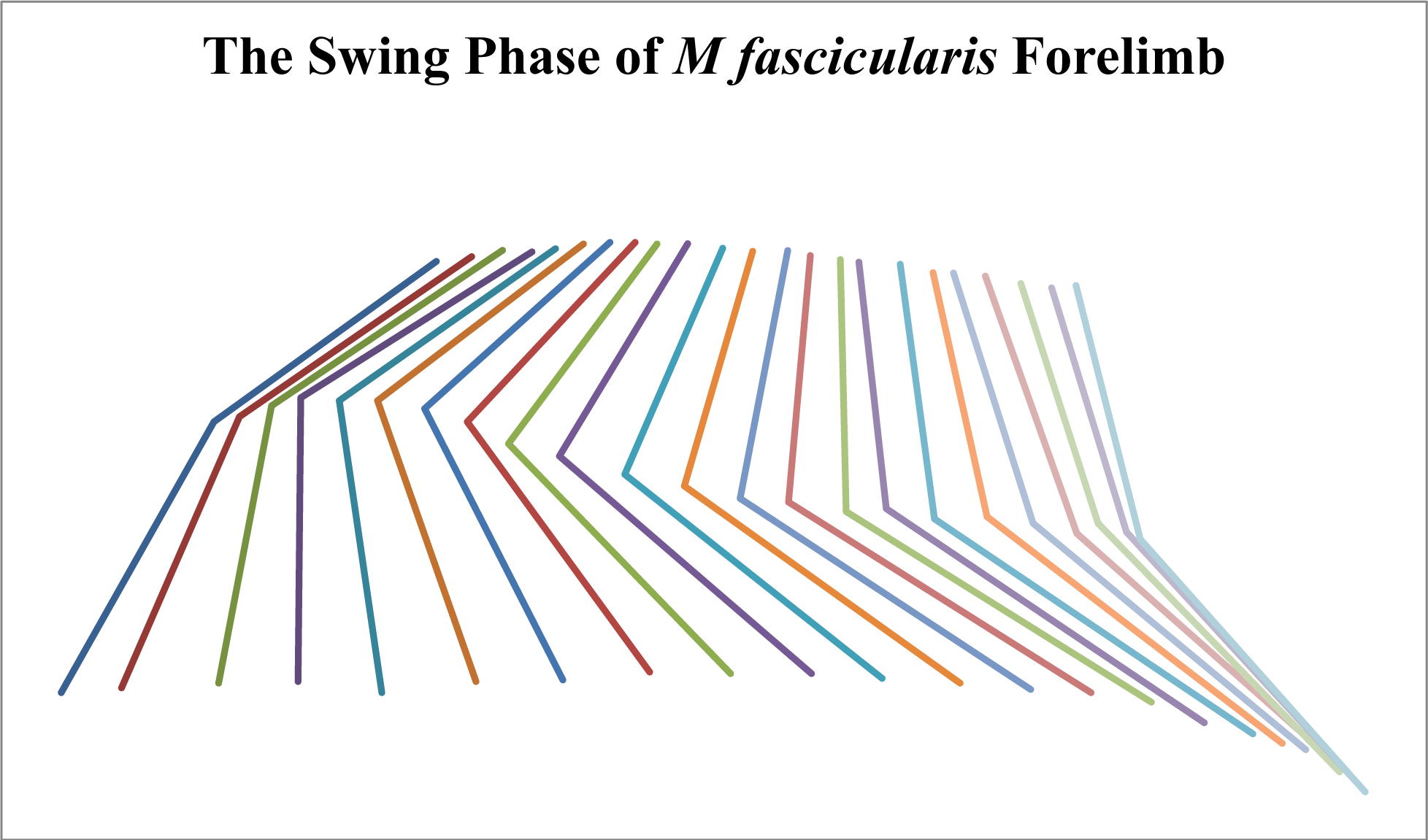

**Figure 10:**
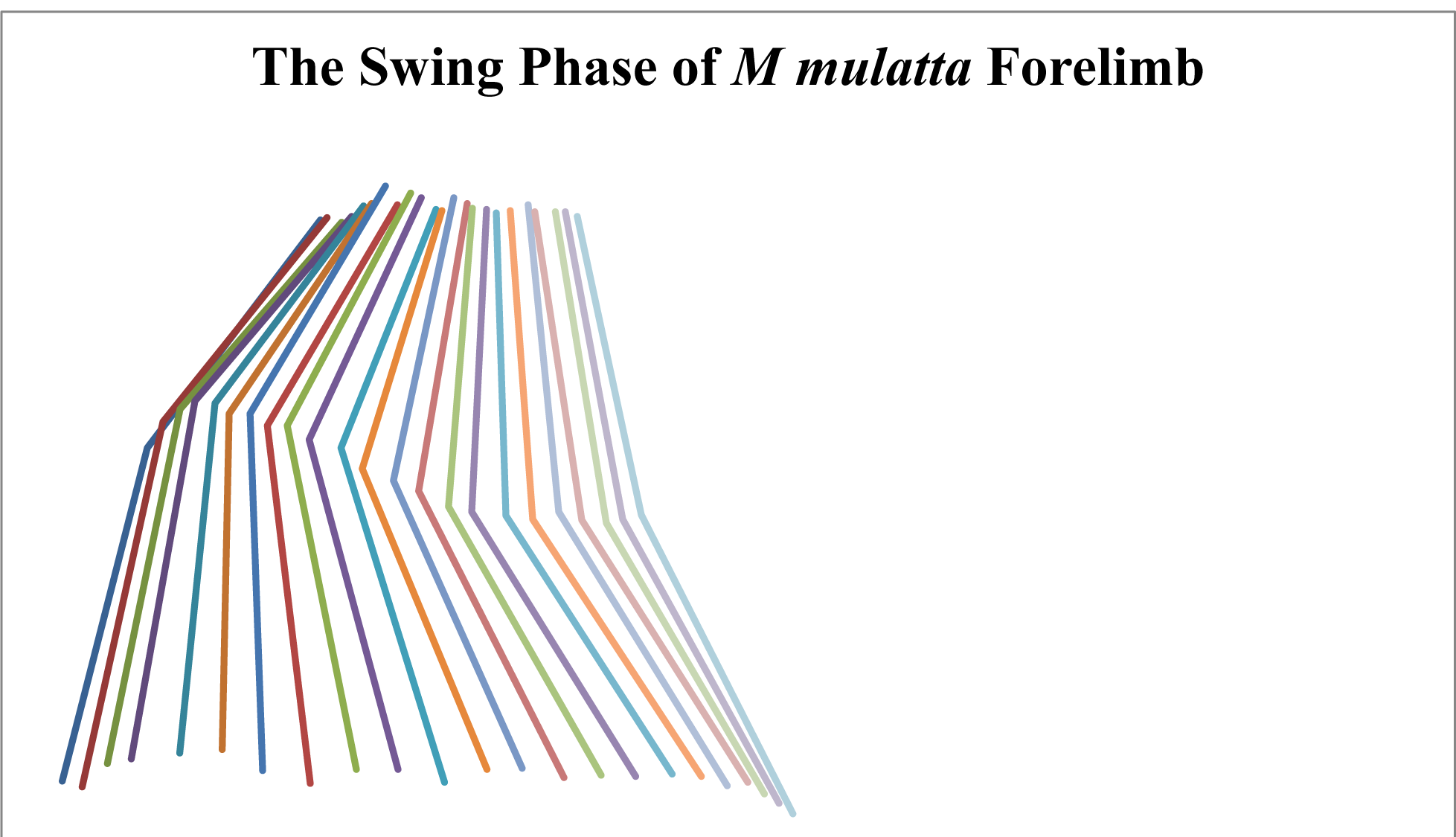

**Figure 11:**
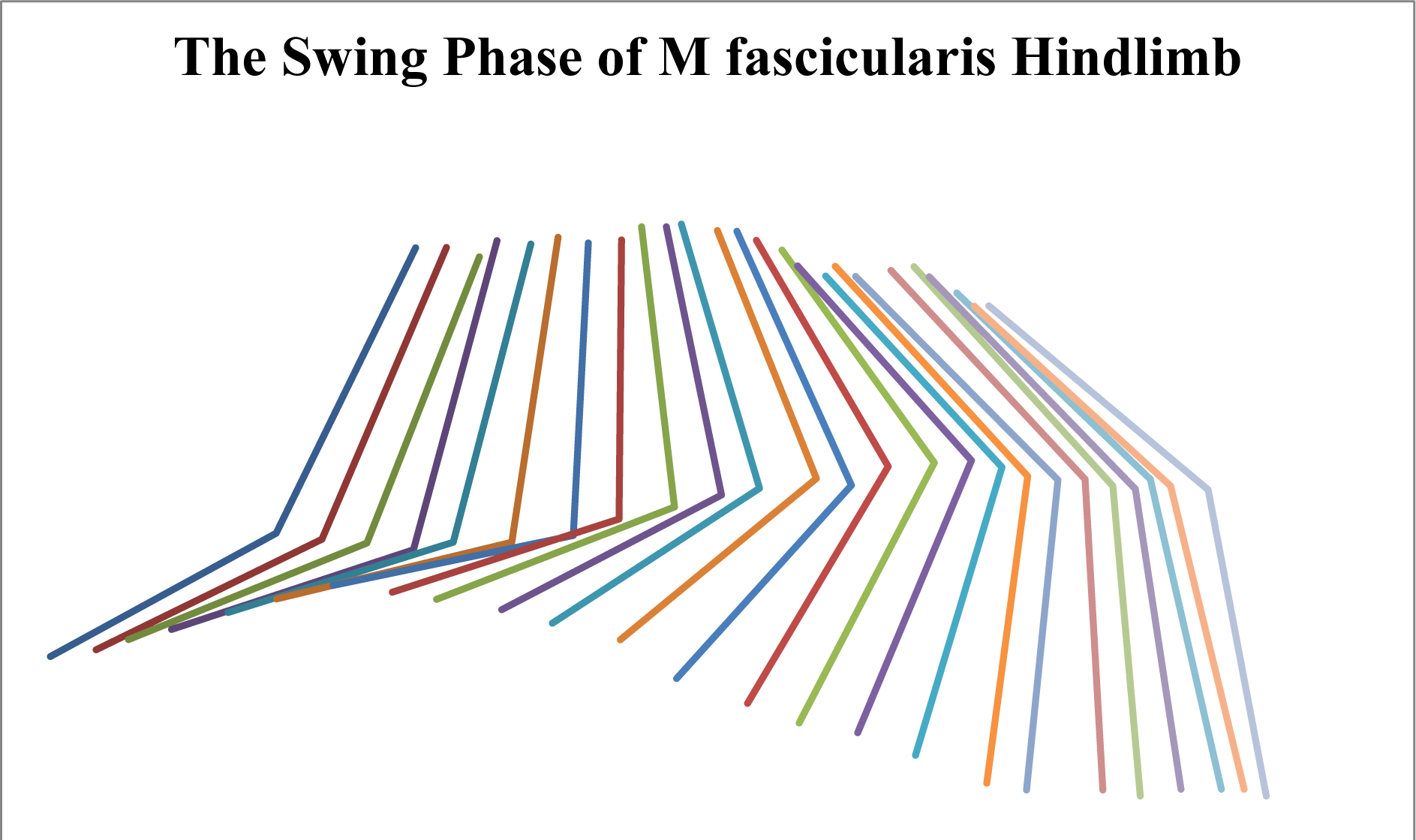

**Figure 12:**
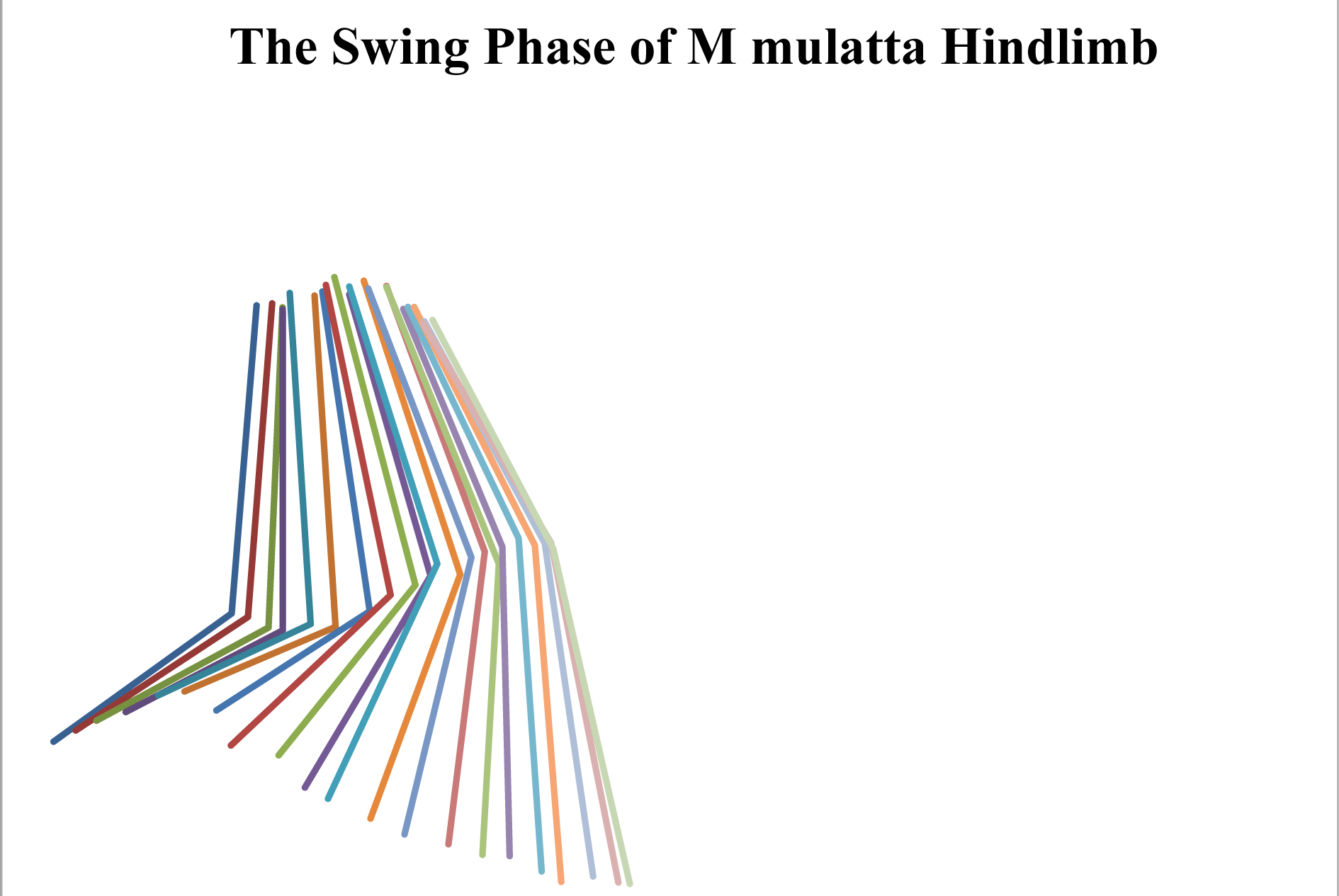

## DISCUSSION

### Hypothesis 1: shoulder and hip retraction

The data do not support the prediction that the more arboreal M. fascicularis would have a greater shoulder retraction angle. The shoulder of the more arboreal *M. fascicularis* has a significantly smaller minimum retraction angle during swing phase than the *M. mulatta*, but does not have a significantly larger maximum retraction angle. Since maximum shoulder retraction occurred shortly after lift-off (Figure 5), it could be due to the animal’s need to clear the substrate so it does not drag its limb at the beginning of swing phase, is unlikely the case because all data on both individuals were collected on similar level terrestrial substrates. It has been shown that the *M. fascicularis* and *M. mulatta* have similar hip retraction angles at lift-off (Schmitt, 1995). A smaller retraction angle means that the humerus reaches a point closer to the center of the body. Also, we would expect the more arboreal *M. fascicularis* to have less of a retracted shoulder, since when walking on thin branches it has the option to swing the limb next to or possibly below the branch, whereas a more terrestrial animal would not have this option. Therefore, it is possible that the increased shoulder retraction serves another purpose. By increasing the retraction of the shoulder the entire limb is brought closer to the body, and more importantly, this change in limb conformation contributes to a movement of the limb center of gravity closer to the body (Figure 9).

The data do not support the idea that during swing phase the more arboreal *M. fascicularis* has a larger minimum hip retraction angle or a larger hip excursion angle than *M. mulatta* as there was no significant difference between the two individuals. Maximum and minimum hip retraction occurred at the beginning and end of swing phase, respectively (Figure 7). It is interesting to note that maximum hip retraction occurred at the initiation of swing phase and therefore did further retract after lift-off. Since the additional retraction was seen in the forelimb and not the hindlimb, it supports the idea that the limb retraction peak is used to clear the substrate, since if this was the case, the additional retraction should be seen in both the fore and hind limb.

### Hypothesis 2: elbow and knee flexion

The data support the hypothesis that the *M. fascicularis* has significantly greater elbow and flexion during swing phase, than the *M. mulatta*. For reasons similar to those mentioned earlier, this difference is unlikely due to a need to clear the substrate. It is most interesting to note the degree of elbow flexion at each stage of swing phase (Figure 6). When comparing the elbow flexion of the *M. fascicularis* and that of the *M. mulatta* it is seems that the maximum elbow flexion occurs earlier during swing phase in the *M. fascicularis*. In the *M. fascicularis*, maximum elbow flexion occurred before mid swing phase, whereas in the *M. mulatta* it occurred approximately at mid swing phase. This suggests that in the *M. fascicularis*, elbow flexion contributes to a limb conformation change that affects the motion and mechanics of swing phase. Since increased elbow flexion bring the middle and distal segments of the limb closer to the shoulder, the point of rotation, it will lead to a more proximal limb center of gravity relative to the limb center of gravity at lift-off. In addition, change in the angle over time when approaching touch-down is seems to be slightly different between the two individuals.

### Preparatory elbow flexion during late swing phase

The elbow angle of the *M. fascicularis* seems have an additional period of flexion right before stance phase. This preparatory muscle action is seen in studies where EMG results depict the pectoralis major and latissimus dorsi muscles showing constant activity from swing phase to stance phase in chimpanzees (Larson, 1998). Marsh et al (2004) also note that some stance phase start activity during late swing phase for guinea fowl. Other studies on the guinea fowl show knee and thigh flexor muscle activity right before swing phase, as well as knee, thigh and foot extensor muscle activity right before stance phase (Gatesy, 1999). This activity likely prepares the limb for stance phase, stiffening it to allow greater mechanical energy conservation. Hip extensor muscles showed activity during late swing phase for a primate sample in Larson and Stern (2009), but no comparative forelimb data exists for *Macaca*.

The findings suggest that further study deserves a strong consideration. These data are supported by the elbow flexion slope approaching zero in the *M. fascicularis* at the end of swing phase. In terms of limb conformation, this means that after maximum flexion, the elbow begins to extend, then a maximum increase in flexion angle occurs *before* the end of swing phase. The increase in flexion angle of the *M. mulatta* seems to stay constant *until* touch-down. It is important for primates to decrease vertical peak substrate reaction forces during stance phase due to their invasion of the small branch niche (Schmitt, 1999; Rollinson and Martin, 1981). Since the *M. fascicularis* has been shown occupy a more arboreal niche (Kurland, 1973) the elbow flexion muscle activity at late swing phase may act as a mechanism to decrease peak vertical substrate reaction forces during stance phase. Essentially activity in the flexor muscles of the elbow would decrease the acceleration of the middle and distal segments of the limb towards the substrate, allowing for a softer impact. This can also be interpreted as a tentative or hesitant motion, which is understandable if the forelimb of arboreal primates must touch down on often unfamiliar or unsteady branches (Rollinson and Martin, 1981).

The data do support the hypothesis that maximum knee flexion in the *M. fascicularis* is greater than the *M. mulatta*. I am hesitant to speculate on the implications of a more flexed knee joint since neither minimum knee flexion or knee flexion angular excursion are significantly different. This suggests that the significant difference in knee flexion may be due to a small sample size.

### Forelimb and hindlimb limb conformation

By looking at the data on forelimb and hindlimb conformation during swing phase I have noticed some interesting comparisons. Since the hindlimb joint angles during swing phase across the two species do not significantly differ, it lends us to believe that in some way the hindlimb function is similar both for arboreal and terrestrial primates. Using the idea of preparatory muscle action described above for the forelimb, it is possible that this sort of action is not necessary for the hindlimb of the *M. fascicularis*.

Primate diagonal sequence gait places the ensuing hindlimb directly next to the previously placed forelimb on the same side (Rollinson and Martin, 1981). Since the forelimb should have already established a secure grip, there would be no reason for the hindlimb to be tentative with its approach. Larson and Stern (2009) actually showed hip extensor activity late in swing phase which not only supports the idea that primates actively shift their weight to their hindlimb (Reynolds, 1985a), but that preparatory muscles of the hindlimb are less worried about increased vertical substrate reaction forces than those of the forelimb.

### Hypothesis 3: limb center of gravity fluctuations

The data from this study support the hypothesis that center of gravity fluctuation, due to limb conformation changes throughout swing phase, are significantly larger in *M. fascicularis* than in *M. mulatta*. It is apparent that the larger vertical change in forelimb center of gravity in the *M. fascicularis* is due to increased retraction at the shoulder and flexion at the elbow during swing phase. This relationship is less apparent in fore the hindlimb, but still requires further study.

### Limb conformation and center of gravity changes

Changes in limb conformation will surely change the location of the limb’s center of gravity throughout swing phase. A more flexed limb conformation during swing phase and its resulting changes to limb center of gravity has significant implications as an adaptation for arboreal locomotion. In effect a more flexed limb conformation allows the animal to have longer limbs with more distal muscle mass, while maintaining a relatively proximal limb center of gravity during swing phase. It is suggested that more arboreal primates have a more distal limb center of mass (Preuschoft and Gunther, 1994).

Although the data on *M. fascicularis* limb center of mass do not support this (Table 2), the data were collected from only one individual. Preuschoft and Gunther (1994) contribute greater distal limb segment mass to an essential arboreal adaptation for greater grasping ability. If this is true, since arboreal primates also show a greater degree of limb flexion during swing phase, their limb center of gravity will reach a higher peak. The further the center of mass is from the most proximal joint, the larger the distance will be that it travels for a given change in angle, conversely, the closer the center of mass is to this point of rotation the shorter the distance will be that it travels for a given change in joint angle. Therefore, arboreal primates with a more distal limb center of mass can change the position of their limb center of gravity to a greater degree. The data support the notion that this is in fact what they do.

**Table 2:** Limb segment masses and dimensionless center of mass (dCM) for the two species compared. Data for M fascicularis was taken from the study at the University of Illionois Aepartment of Anthropology on a single subject cadaver. Data for M mulatta was taken from Vilensky (1979).

| **Segment** | **Mass (g)** | **dCM (% segment length)** |
| --- | --- | --- |
| *M fascicularis* |  |  |
| Arm | 134.4 | 0.41 |
| Forearm and hand | 95.1 | 0.43 |
| Thigh | 231.5 | 0.45 |
| Leg and Foot | 142.4 | 0.36 |
| *M mulatta* |  |  |
| Arm | 218.1 | 0.48 |
| Forearm and hand | 163 | 0.59 |
| Thigh | 363 | 0.51 |
| Leg and foot | 243.7 | 0.63 |

**Table 3:** Comparison of variables between *M fascicularis* and *M mulatta* subjects, showing significant p-values. For clarity, hip and shoulder retraction angle were treated as absolute angles and not as amount of protraction or amount of retraction as mentioned above. For elbow and knee flexion, larger or maximum flexion angles represent a more flexed and less extended joint, and smaller or minimum flexion angles represent a less flexed joint and more extended joint.

| **Variable** | **p-value** |
| --- | --- |
| **Hip retraction angle** |  |
| Max | ns |
| Min | ns |
| Angular excursion | ns |
| **Shoulder retraction angle** |  |
| Max | ns |
| Min | 0.016649 |
| Angular excursion | 0.040628 |
| **Knee flexion angle** |  |
| Max | 0.038561 |
| Min | ns |
| Angular excursion | ns |
| **Elbow flexion angle** |  |
| Max | < 0.001 |
| Min | ns |
| Angular excursion | < 0.001 |
| **Center of gravity** |  |
| Forelimb Cg max | 0.012186 |
| Forelimb Cg min | ns |
| Forelimb Cg range | 0.021572 |
| Hindlimb Cg max | 0.007546 |
| Hindlimb Cg min | 0.013711 |
| Hindlimb Cg range | ns |

**Table 4:** Significance test results for manipulated limb segment center of mass data.

| Variable | p-value |
| --- | --- |
| Forelimb Cg* Max | 0.012186 |
| Hindlimb Cg* Max | 0.007546 |

#### Limb center of gravity: implications for mechanical energy conservation

This study supports the idea that for the *M. fascicularis* compared to the *M. mulatta*, the forelimb and hindlimb limb center of gravity are brought significantly closer to the body at the beginning of swing phase. Since the location of the limb center of gravity is directly related to its moment of inertia and Natural Pendular Period (NPP) (Myers and Steudel, 1997) the changes in limb center of gravity in the *M. fascicularis* surely affect its moment of inertia during swing phase. Raichlen (2004) found that at maximum flexion, the NPP of his baboon sample was reduced by 12%. Myers and Steudel (1997) found that it changed the NPP of their canine sample by 7%. Since the change in the baboon sample was greater than the canine sample, it supports the idea that primate limb conformation and limb center of gravity have a greater affect on the inertial properties of the limb than do those of the non-primates.

Although I am unable to say for certain, I would assume that the difference in limb center of gravity in those two samples was due a more distal limb center of mass, and a greater range of joint flexion in the primate (baboon) sample. Rodman (1979) suggests that greyhounds should have lighter distal limb segments than primates due to a greater proportion of time spent walking on the ground. In addition, the hands and feet of *M. fascicularis* are heavier than those of *M. nemestrina*, a more terrestrial macaque species (Rodman, 1979). If the energetics of progression become increasingly important percent arboreality increases (Rodman, 1979) then it is likely that this change in limb center of gravity does in fact serve as a mechanism on mechanical energy conservation for more arboreal primate species.

Since a more flexed limb conformation during swing phase causes a decrease in the limb moment of inertia (Raichlen, 2004; Crompton, Li, Alexander, Wang, and Gunther, 1996), the change in inertia may actually contribute to the swinging of the limb. Changing the moment of inertia of the limbs during swing phase resembles the motion of an ice skater (Crompton et al, 1996). If the skater brings their arms closer to their body when spinning, their moment of inertia decreases and they spin faster without doing any additional work. This sample theory can be applied to initiation of the swing phase of the limb. By decreasing the moment of inertia of the limb, the limb can be accelerated through swing phase with out adding addition energy apart from the initial energy used to flex the limb. The reverse is true for extending the limb center of gravity away from the point of rotation. Here the moment of inertia would increase causing the limb swing to slow down at the end of swing phase. If in these scenarios the timing and change in magnitude of flexion and extension were coordinated in such a way that they no additional energy was needed to swing the limb relative to the body, energy would be conserved to a greater degree. Since Marsh et al (2004) found that swing phase may consume up to 26% of the total energy during each step a more efficient swing phase would serve as a good way to conserve total mechanical energy.

Primates use a relatively more compliant gait compared to non-primates (Schmitt, 1999), which is due to their occupation of the arboreal environment (Rollinson and Martin, 1981). Stiff limbs are more effective in mechanical energy conservation during stance phase due to the larger vertical oscillations of the body’s center of mass they create, yet they also cause larger peak vertical substrate reaction forces (Schmitt, 1999), so large to and midsized arboreal primates have adopted a more compliant limb posture. Since the compliant limb posture during stance phase does not conserve mechanical energy as effectively as a stiff limb posture, these primates likely have found another mechanism for mechanical energy conservation. It is suggested by the data that the more arboreal primate species *M. fascicularis* uses limb conformation change to adjust its limb moment of inertia in order to conserve mechanical energy during swing phase. It is not necessary for the more terrestrial *M. mulatta* to adopt this method of limb swinging because it is able to maintain stiff limbs during stance phase, and it does not have to maneuver on thin branches as often as the *M. fascicularis*.

## CONCLUSION

In this study I suggest how locomotor variation between two closely related primate species, *M. mulatta* and *M. fascicularis*, can be explained by implications of mechanical energy conservation. I show that the mechanics of swing phase for these two species have important effects on their limb center of gravity, although they may have similar limb segment mass distributions. This study suggests that not only is mechanical energy conservation during swing phase important in mammalian locomotion, but that its complexity deserves further consideration and analysis.

## ACKNOWLEDGMENTS

I would like to thank John Polk from the University of Illinois Department of Anthropology for allowing me to visit his lab and collect crucial data for my research. I would also like to thank the Duke Undergraduate Research Support for providing the funding necessary for all aspects of this study. I would like to thank my major advisor, Leslie Digby, for the encouragement that gave me the confidence to take on this research. Finally, I would like to give additional thanks to Dan Schmitt and my mentor Tracy Kivell for their guidance, support, and insightful discussions throughout the duration of my research.

---

**APPENDIX 1:**
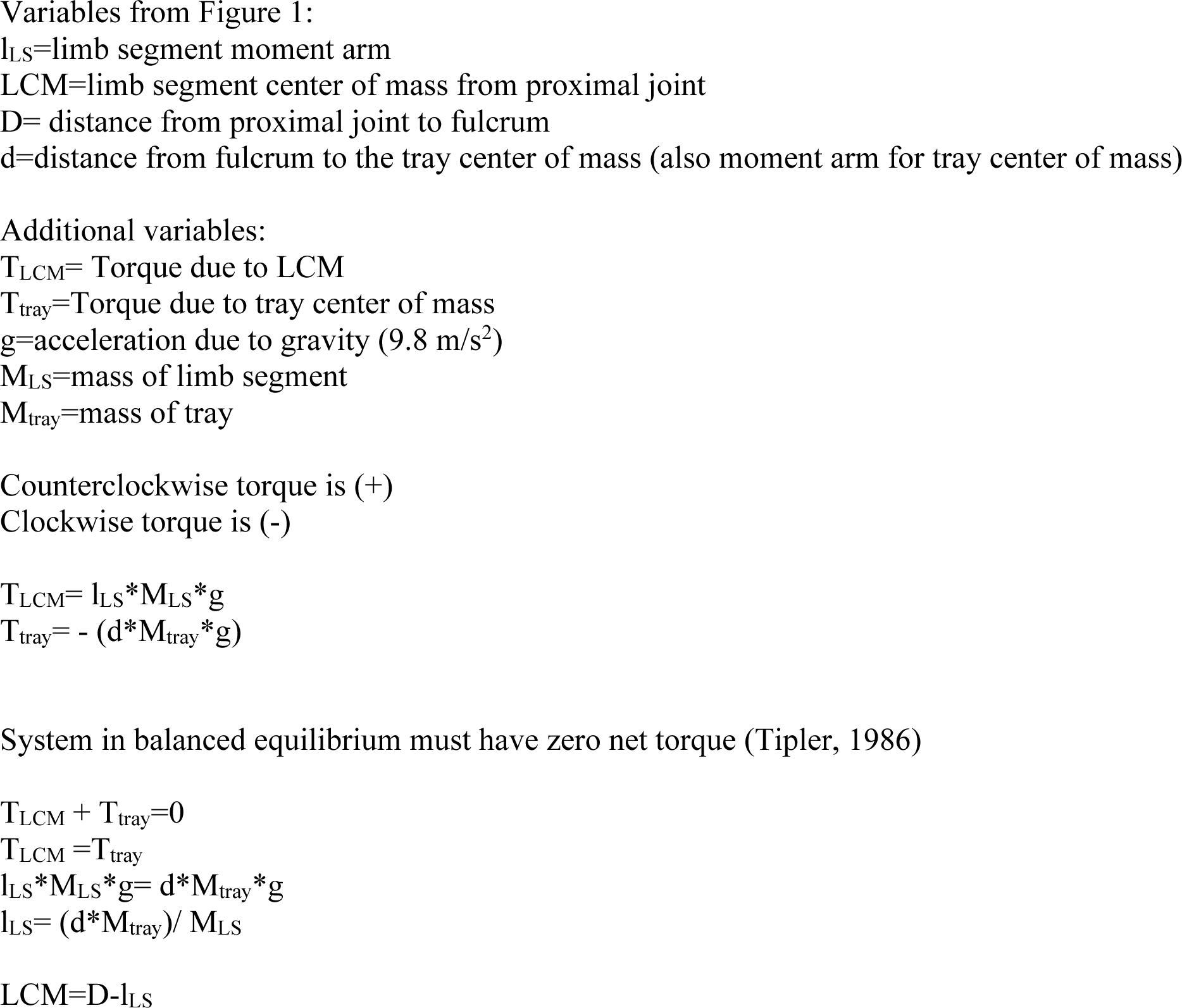
Limb segment center of mass using summation of torques

**APPENDIX 2:**
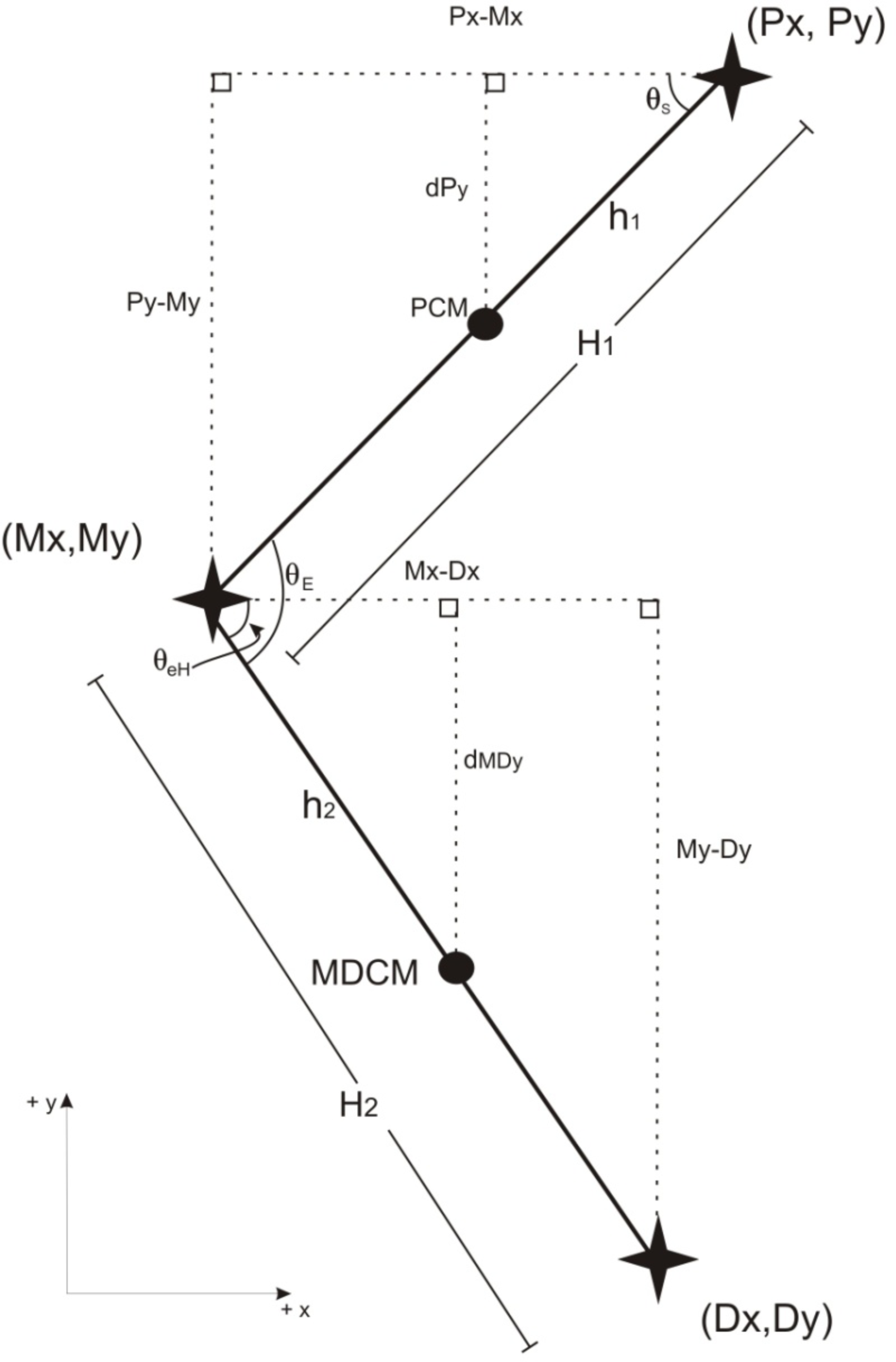
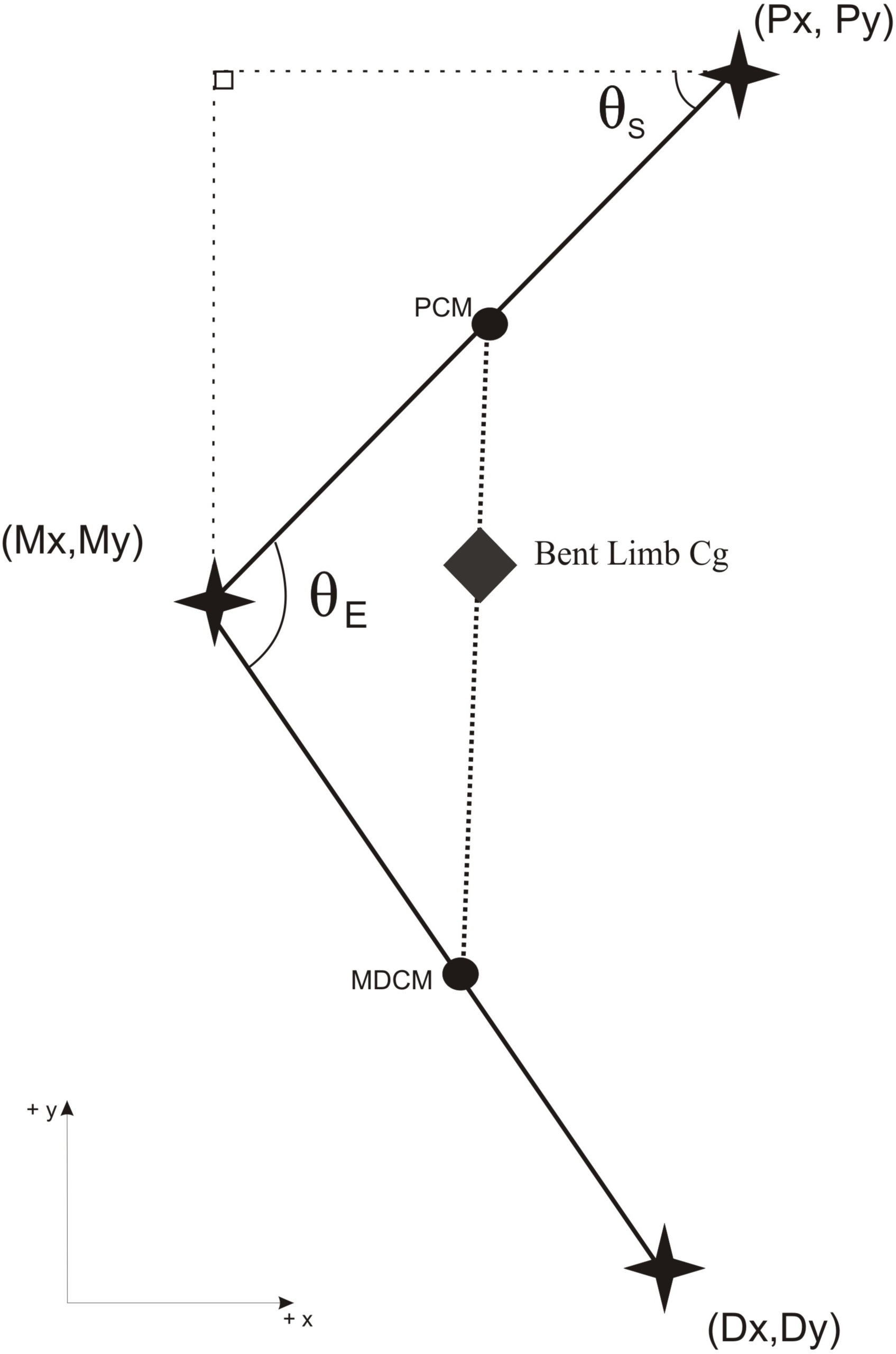
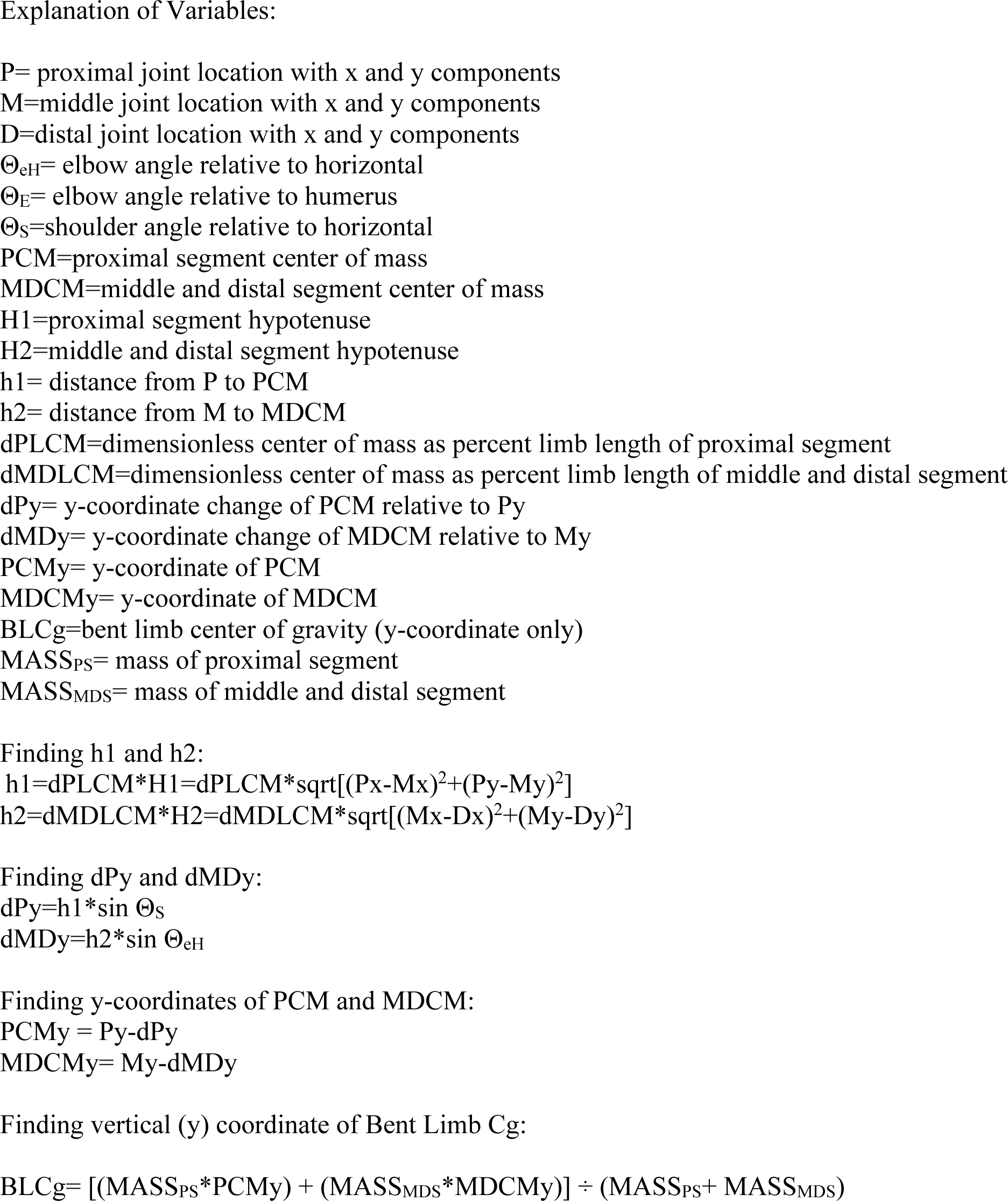
Finding vertical component (y) of bent limb center of gravity (forelimb example)

## Footnotes

* Center of gravity and center of mass are interchangeable, and both describe one point where the summation of parallel weight vectors produce zero torque (Tipler, 1986). Although these phrases are essentially the same, we will use center of mass to describe the location of a point on the limb itself somewhere between the tip of the most distal segment and the most proximal end, as if the limb represented a straight rigid segment. When referring to center of gravity, we mean the point in space between the proximal segment center of mass and the middle segment center of mass when the limb is going through conformational change. Therefore the center of gravity may not lie on the limb itself, as would be the case if the limb was bent.

1 Please note for this study, I have defined retraction angle as the angle created by the limb and the horizontal as shown above. Retraction will be expressed as an anterior/posterior movement toward the center of the body and protraction will be defined as an anterior/posterior movement away from the body. During swing phase the forelimb actually begins retracted and finishes protracted and the hindlimb begins protracted and finishes retracted, since at 90 degrees the naming convention changes. To avoid confusion, I will refer to all hip and shoulder angles as retraction angles. For reference, a retraction angle of θ is equivalent to a protraction angle of 180 – θ.

